# Impact of a Patient-Derived Hepatitis C Viral RNA Genome with a Mutated MicroRNA Binding Site

**DOI:** 10.1101/469841

**Authors:** Miguel Mata, Steven Neben, Karim Majzoub, Jan Carette, Peter Sarnow

## Abstract

Hepatitis C virus (HCV) depends on liver-specific microRNA miR-122 for efficient viral RNA amplification in liver cells. This microRNA interacts with two different conserved sites at the very 5’ end of the viral RNA, enhancing miR-122 stability and promoting replication of the viral RNA. Treatment of HCV patients with oligonucleotides that sequester mir-122 resulted in profound loss of viral RNA in phase II clinical trials. However, some patients accumulated in their sera a viral RNA genome that contained a single cytidine to uridine mutation at the third nucleotide from the 5’ genomic end. It is shown here that this C3U variant indeed displayed higher rates of replication than that of wild-type HCV when miR-122 abundance is low in liver cells. However, when miR-122 abundance is high, binding of miR-122 to site 1, most proximal to the 5’ end in the C3U variant RNA, is impaired without disrupting the binding of miR-122 to site 2. As a result, C3U RNA displays a much lower rate of replication than wild-type mRNA when miR-122 abundance is high in the liver. These findings suggest that sequestration of miR-122 leads to a resistance-associated mutation that has only been observed in treated patients so far, and raises the question about the function of the C3U variant in the peripheral blood.

**Author Summary:** With the advent of potent direct-acting antivirals (DAA), hepatitis C virus (HCV) can now be eliminated from the majority of patients, using multidrug therapy with DAAs. However, such DAAs are not available for the treatment of most RNA virus infections. The main problem is the high error rate by which RNA-dependent RNA polymerases copy viral RNA genomes, allowing the selection of mutations that are resistant to DAAs. Thus, targeting host-encoded functions that are essential for growth of the virus but not for the host cell offer promising, novel approaches. HCV needs host-encoded microRNA miR-122 for its viral RNA replication in the liver, and depletion of miR-122 in HCV patients results in loss of viral RNA. This study shows that a single-nucleotide mutation in HCV allows viral RNA amplification when miR-122 abundances are low, concomitant with changes in its tropism.

## Introduction

Many cell- and virus-encoded microRNAs (miRNAs) regulate the expression of mRNAs by binding to the 3’ noncoding regions of target mRNAs. The binding is facilitated by an RNA-induced silencing complex (RISC) that mediates base-pair interactions between nucleotides two through seven in the microRNA (seed sequences) and their complementary sites in the target mRNA (seed-match sequences). This targeting event inhibits the translation of the mRNA. In addition, deadenylation at the 3’ of the mRNA, followed by decapping and 5’ to 3’ degradation of the mRNA greatly increases its turnover [1, 2].

The growth of hepatitis C virus (HCV), a member of the flaviviridae, is dependent on the most abundant miRNA in the liver, miR-122 [3]. In the liver, miR-122 is known to be crucial for upregulation of cholesterol metabolism [4, 5]. In the HCV genome, we discovered two binding sites for miR-122 at the 5’ proximal end of the viral RNA [3]. Occupancy of both sites by miR-122 is required for the maintenance of viral RNA abundance in infected liver cells [3, 6-8]. Loss of HCV RNA abundance could be observed when HCV-infected cells were treated with modified oligonucleotides that have base-pair complementarity to miR-122 (miR-122 anti-miRs) [3]. HCV sequences termed site 1 and site 2 (Fig 6A) are seed-match sequences for miR-122 and are both absolutely conserved among all genotypes of HCV and present in all HCV gene sequences from patients deposited in gene banks. Deleterious effects of mutations in either miR-122 binding site 1 or site 2 on HCV RNA accumulation can be rescued by co-transfection of mimetic miR-122 duplexes that targeted the mutated HCV genomes [3, 6] [7]. Thus, HCV subverts host miR-122 to increase its expression in liver cells. It is envisaged that HCV RNA genomes have evolved to bind the highly abundant liver-specific miR-122 [9] to guarantee persistence in the liver over many years. Mechanistically, miR-122 has been shown to protect the 5’ ppp-containing HCV genome from the action of 5’ RNA triphosphatase DUSP11 [10, 11] and subsequent degradation by 5’ RNA exonucleases XRN1 [12] and XRN2 [13]. Further evidence suggests that miR-122 also participates in the switch of viral RNAs from the translation to the replication phase in the viral life cycle by displacing of RNA binding proteins that enhance viral mRNA translation [14].

Clinical applications of miR-122 anti-miRs first showed that sequestration of miR-122 in mice [4, 5] and in non-human primates [15] lowered plasma and liver cholesterol abundance without any obvious adverse effects on liver function. Subsequently, Lanford and colleagues [16] tested the effects of sequestration of miR-122 after intravenous administration of unformulated locked nucleic acids (LNAs)-containing miR-122 anti-miRs in HCV-infected chimpanzees. Administration of LNAs caused a 500-fold reduction of viral titer in both serum and liver that persisted for several weeks. Encouraged by these results, independent studies evaluated the efficacy of two different miR-122 anti-miRs, miravirsen [17] and RG101 [18], in patients with chronic HCV genotype 1 infections. Treated patients showed a 10-1000 fold reduction in viral serum titers. While the majority of the patients cleared the infection, several subjects in both studies experienced a virologic rebound several weeks after anti-miR treatment [18, 19]. Sequence analysis of viral RNAs obtained from serum of several of those patients revealed a resistance-associated substitution of a uridine for a cytidine nucleotide 3 (C3U) [17, 18]. The goal of the present study was to examine whether the C3U mutation truly confers resistance to miR-122 anti-miRs and to determine the mechanism of any such escape, to provide a basis for investigation whether these variants will compromise clinical treatments.

## Results

### C3U HCV RNA replicates with higher efficiency than wild-type HCV RNA during miR-122 sequestration

Sera from five out of six HCV patients whose viral titers rebounded following miR-122 anti-miR treatment contained HCV genomes with a C3U mutation. One patient contained a C3U/C27U double mutation [18]. These findings suggested that the C3U RNA variant may replicate with higher efficiency than wild-type RNA in liver cells when miR-122 is sequestrated, and thus represent a drug-resistant variant.

To test this hypothesis, in vitro-synthesized wild-type and C3U *Gaussia* luciferase-expressing H77S.3/GLuc viral RNAs (Fig 1A) [20] were transfected into Huh7.5 cells that had previously been treated with non-122 targeting control anti-miR-106b-LNA, anti-miR-122-LNA miravirsen [19], or anti-miR-122 RG1649 (the active metabolite of RG101) [18], and HCV RNA replication was monitored. Figure 1B shows that treatment of HCV RNA-transfected cells with miR-106b-LNA had no effect on wild type HCV RNA translation or replication, as reflected by the accumulation of luciferase encoded in the viral genome. However, wild-type HCV RNA accumulation was reduced relative to C3U HCV in miR-122 anti-miR-treated cells after 24 and 48 hours. These findings argue that reduced abundance of miR-122 is less inhibitory to the growth of C3U HCV, thus rationalizing the emergence of the C3U genome in anti-miR-treated patients.

**Fig 1.**
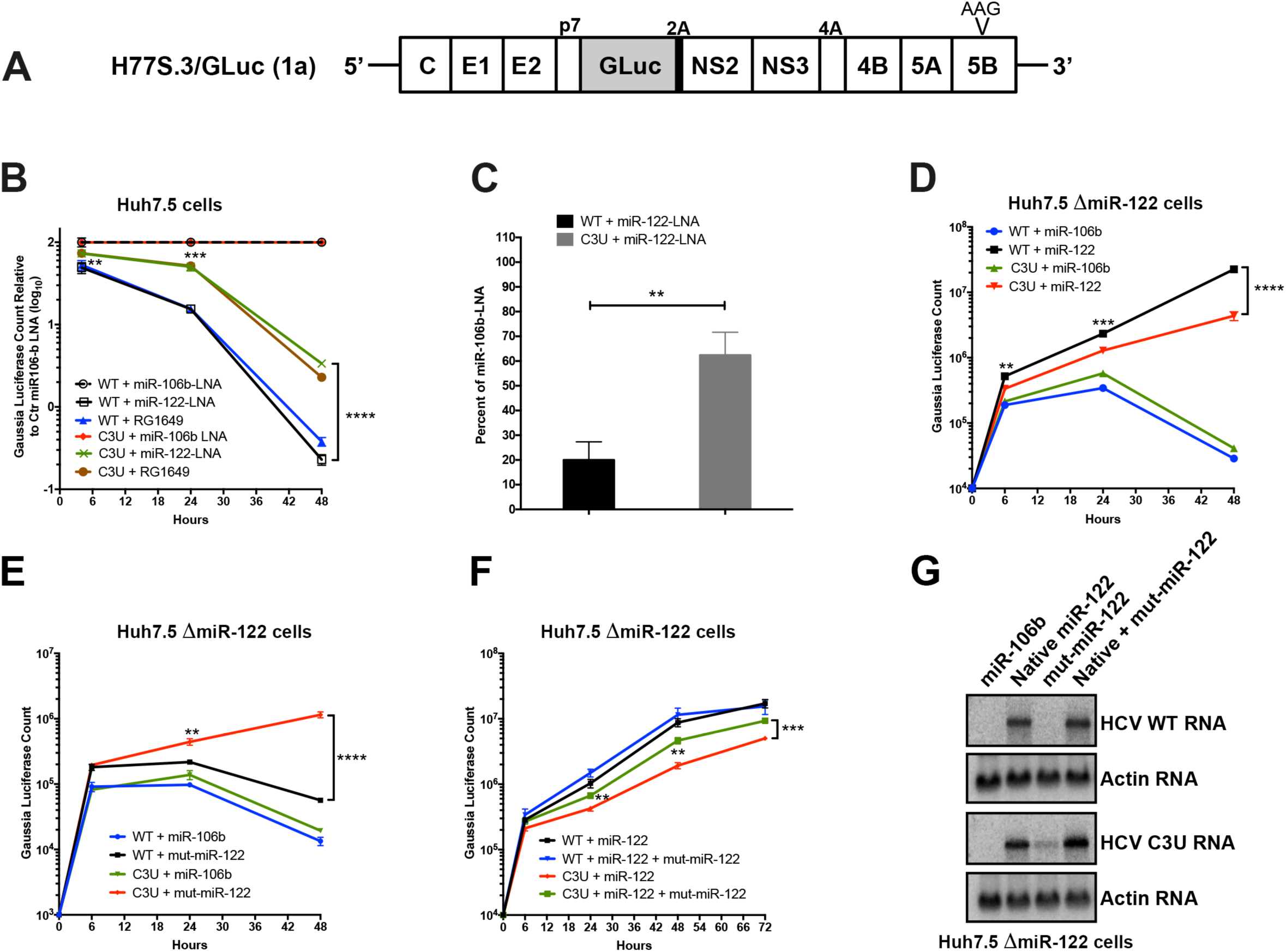
Effects of sequestration or loss of miR-122 on wild-type and C3U variant RNA abundances and replication. (A) Top: Structure of the infectious H77.S3/GLuc construct, containing Gaussia luciferase (GLuc) and foot-and-mouth disease virus 2A autoprotease 2A between the p7 and NS2 sequence [20]. Replication defective mutation (AAG) in the NS5B viral polymerase gene is marked. (B) Effects of miR-122 sequestration using miR-106 LNA, miR-122-LNA or RG1649 anti-miRs. Viral RNA replication was measured by luciferase production at the indicated time points. Data is presented relative to the luciferase expression of negative control miR-106b-LNA treatment. (C) Rate of replication following miR-122 sequestration. Cells were treated as in (B) and 24 hours post RNA transfection, RNA was labeled with 200 μM 5-ethynyl uridine (EU) for twenty-four hours, conjugated to biotin and subsequently isolated with streptavidin beads. RNA replication rates were determined by qRT-PCR. Data is presented as percent labeling relative to of miR-106b-LNA control treated samples (**p<0.0034). (D-F) HCV RNA replication in ΔmiR-122 Huh7 transfected cells. Cells were pretreated with miR-106b, native miR-122, mut-miR-122 alone or in combination one day before and one day after HCV RNA transfection. Supernatants were collected at the indicated time points and GLuc activity was determined. The data are representative of 3 independent replicates (**p<0.005, ***p<0.001 ****p<0.0001). (G) HCV and actin RNA abundances, monitored by Northern blot analysis, at 72 hours post RNA transfection.

To determine whether the C3U mutation acts by increasing the abundance of HCV RNA following miravirsen-mediated sequestration of miR-122, the accumulation of newly synthesized viral RNA was measured by labeling with 5-ethynyl uridine (5-EU), an analog of uridine. Total RNA was isolated and conjugated to biotin in a copper-catalyzed reaction. Newly synthesized RNA was subsequently captured using magnetic streptavidin beads and quantitated by qRT-PCR using HCV-specific primers. While the abundance of wild-type RNA synthesis was reduced by ~80% in miravirsen-treated samples, only a 30% reduction was observed in C3U variant samples following miR-122 depletion compared to control miR-106b-LNA treatment (Fig 1C).

To determine whether the binding of miR-122 molecules to the C3U HCV genome plays any role at all in its replication, infectious RNAs were transfected into miR-122 knock-out Huh7.5 cells [21] that were pre-transfected with control duplex miR-106b or native miR-122 duplexes. Exogenously added control duplex failed to rescue the greatly reduced abundance of either wild-type or C3U RNA. Supplementation of miR-122 duplexes enhanced the amplification of both wild-type and mutant viral RNAs, albeit C3U to a significantly lesser extent than wild-type RNA (Fig 1D). These results argue that C3U RNA requires at least a small amount of miR-122. It is known that miR-122 has different affinities to site 1 and 2 in HCV RNA [22]. Thus, different affinities for miR-122 at site 1 or site 2 in both C3U and wild-type RNAs could explain the observed phenotype.

To test this hypothesis, C3U-complementary duplex miR-122 molecules (referred to as “mut-miR-122”) were employed to monitor C3U RNA replication phenotypes in cells that expressed no wild-type version of miR-122. Again, pre-treatment with miR-106b RNA mimetics did not rescue replication of wild-type or C3U viral RNAs (Fig 1E). As was also expected, supplementation of mut-miR-122 failed to efficiently support wild-type viral RNA accumulation. However, supplementation of C3U with mut-miR-122 efficiently enhanced C3U RNA abundance. That both wild-type and mutant miR-122 can rescue the C3U genome to some extent, but neither as completely as native miR-122 rescues the wild-type genome, suggests that both wild-type and mutant miR-122 might bind to the C3U genome.

To substantiate this finding further, miR-122 knock-out cells were pre-transfected with wild-type and mut-miR-122 duplexes alone or in combination and abundances of transfected infectious RNAs were monitored over time. Co-transfection of both wild-type and mut-miR-122 had minimal additive effect on the accumulation of wild-type HCV RNA compared to transfection of wild-type miR-122 alone (Fig 1F). In sharp contrast, accumulation of the C3U variant was significantly enhanced in the presence of combined wild-type and mut-miR-122 mimetics compared to the presence of either alone (Fig 1F). Analysis of viral infections by Northern blot analyses (Fig. 1G) revealed the same phenotypes. These data show that a single C3U nucleotide substitution in the 5’ noncoding region of the HCV genomic RNA, where miR-122 occupies binding site 1, results in increased resistance to miR-122 sequestration. Because HCV RNA accumulation in a C3U background could be effectively rescued only by a combination of native and mutant-miR-122 mimetics, sites 1 and 2 in C3U RNA are likely occupied by mutant and wild-type miR-122, respectively, and occupancy of both is required for its optimal function. This situation is not likely to be obtained under any circumstance in liver cells, leading us to question whether the C3U mutation might bring about a large fitness cost to viruses that contain it.

### C3U mutation in HCV RNA reduces viral fitness in cultured liver cells

To investigate the impact of the C3U HCV mutation on viral fitness in liver cells when miR-122 abundance is normal, in vitro-synthesized wild-type and C3U H77.S3 infectious RNAs [20] were electroporated into Huh7.5 liver cells [23]. At different times after electroporation, supernatants were collected and extracellular HCV titers were determined by fluorescent focus forming units (FFU) measurements after infection of Huh7.5 cells (Fig 2A). In contrast to wild-type virus-infected cells, which reached a maximum of 10^3^ FFU/ml, virus yield was ten-fold lower in cells infected with C3U HCV virus (Fig 2A). Next, viral spread was examined by infecting naïve Huh7.5 cells with wild-type and mutant viruses. Supernatants were collected three days post-infection and quantified by FFU assays. Similarly, to the data shown in Fig. 2A, viral production (Fig. 2B) and RNA accumulation (Fig. 2C) after these multiple cycles of infection were ten-fold lower in C3U HCV-infected compared to wild-type HCV-infected cells. Thus, the C3U mutation results in a significant reduction in mutant viral RNA and virion abundances at a step that does not involve viral entry.

**Fig 2.**
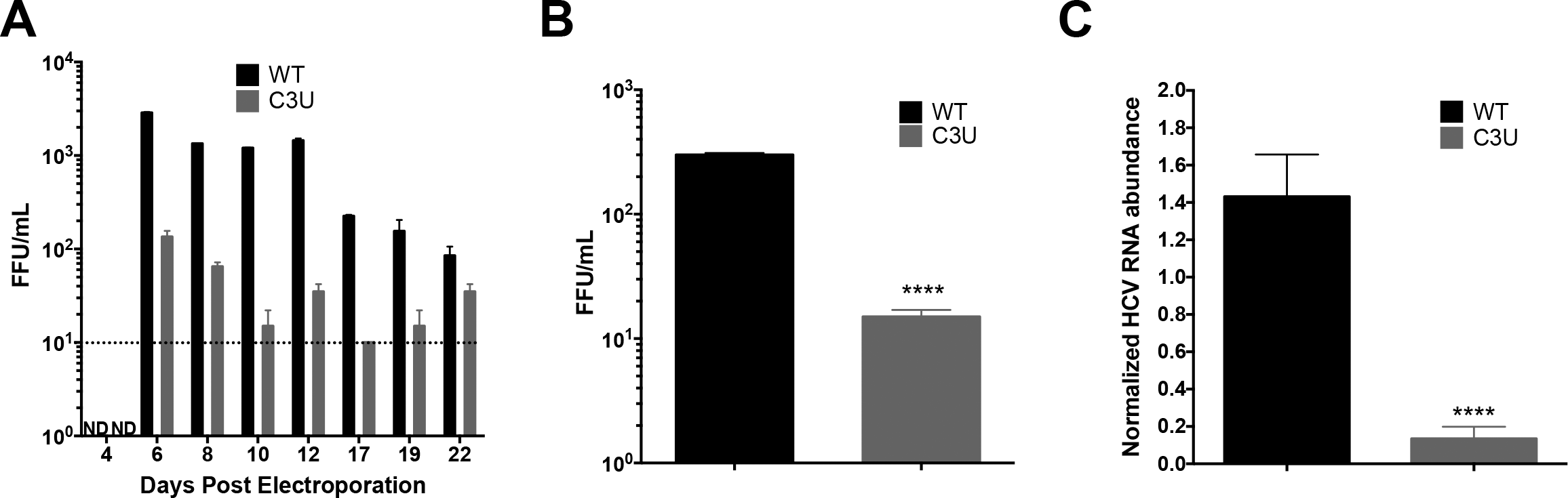
Effects of the C3U single nucleotide substitution on HCV genotype 1a virus production. (A) Extracellular virus production. Huh7.5 cells were electroporated with 10 μg of infectious HCV genotype 1a (H77) RNA that does not (WT) or does contain the C3U mutation. Titers were determined by focus forming assays (FFU). Dotted line indicates threshold of detection. (B) Virus titers of Huh7.5 cells infected with wild-type and C3U H77 variants at a multiplicity of infection (moi) of 0.005 and assayed at 72 hours post infection (**** p<0.0001). (C) RT-qPCR measurement of HCV viral RNA abundance at 5 days post-virus infection at an MOI of 0.005. Data normalized to internal control 18S rRNA levels (**** p<0.0001). Error bars display +/- SD.

### C3U mutation leads to reduced viral RNA replication in liver cells

To measure the effects of the C3U mutation on viral mRNA translation and RNA replication, in vitro-synthesized wild-type and C3U *Gaussia* luciferase-expressing H77S.3/GLuc viral RNAs (Fig. 1A) were transfected into Huh7.5 cells and luciferase activities were examined at different times after transfection. Replication of the C3U viral RNA was ten-fold lower than that of wild-type RNA between 48 and 72 hours after transfection (Fig 3A). Similar to the phenotype observed with luciferase activities, Northern blot analysis revealed a decrease of C3U RNA abundance compared to wild-type viral RNA at three days after RNA transfection (Fig 3B). Both wild-type and mutant RNAs were sensitive to the HCV RNA polymerase NS5B inhibitor sofosbuvir (Fig. 3B), demonstrating that authentic viral RNAs were inspected in the Northern blots and that the C3U variant can be eliminated by sofosbuvir. The effect of the C3U mutation was not genotype-specific, because insertion of the C3U mutation into the J6/JFH1 RLuc genotype 2a infectious background also resulted in significant reduction in RNA replication (Fig S1). Finally, the abundance of viral core protein was also reduced in C3U RNA-transfected cells at 72 hours compared to wild-type RNA (Fig. 3C), suggesting defects in protein synthesis, RNA replication or both in C3U HCV.

**Fig 3.**
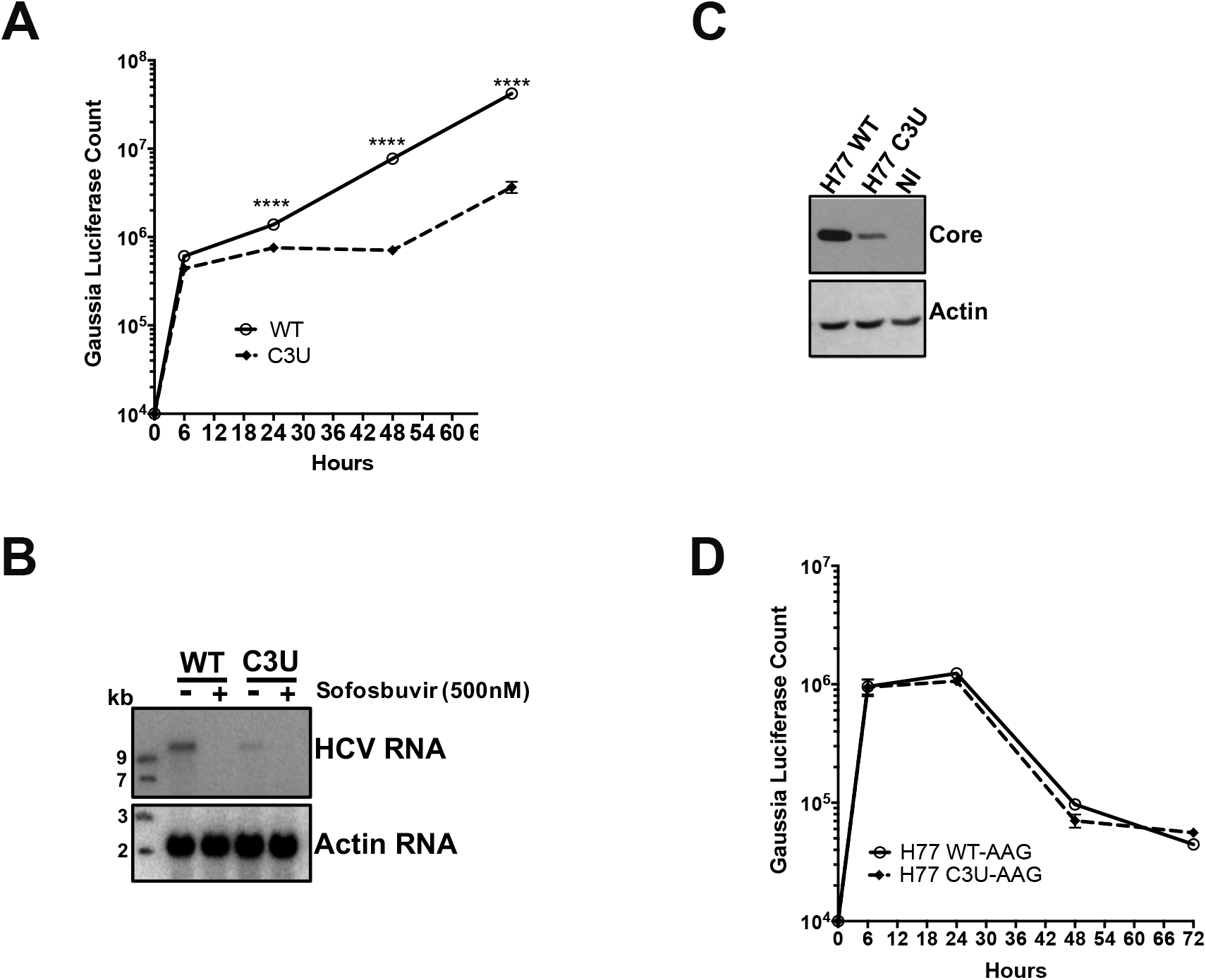
Effects of C3U mutation on HCV RNA replication and translation. (A) HCV RNA replication was monitored by the expression of GLuc secreted into the supernatants of wild-type and C3U RNA transfected Huh7.5 cells. Supernatants were collected at the indicated time points. (B) Wild-type and C3U H77.S3/GLuc RNA abundances three days post-transfection in untreated and cells treated with the NS5B inhibitor sofosbuvir (500nM). (C) Effects of C3U mutation on HCV core protein levels examined by Western blot analysis at three days post-H77.S3/GLuc RNA transfection. (D) Translation of the replication defective (AAG) wild-type and C3U H77.S3/GLuc RNAs at multiple several points post-transfection. The data are representative of 3 independent replicates (****p<0.0001, Student’s t-test).

To examine in detail whether reduced translation or replication contributed to low viral RNA abundances in C3U RNA-transfected cells, we studied the expression of chimeric RNAs, containing GDD-to-AAG mutations in the active site of viral RNA-dependent polymerases NS5B in both wild-type and C3U RNAs. Figure 3D shows that translation of the C3U variant was indistinguishable from wild-type at all time points measured, arguing that the growth defect of C3U HCV is predominantly at the replication step. To examine effects of the C3U mutation on HCV translation by a different approach, polysomal mRNAs were analyzed from HCV RNA-transfected cells after separating cell lysates in sucrose gradients. The distribution of full-length HCV RNA in each individual fraction was analyzed by Northern blot analysis (Fig. S2A). HCV RNA was distributed in polysomal fractions 9 through 13 in both wild-type- and C3U-transfected samples (Fig S2A, B). These data suggest that the observed reduced intracellular RNA abundance of C3U HCV is primarily due to a defect in RNA replication or stability.

### Effects of the C3U substitution on viral RNA stability and nascent viral RNA synthesis in the presence of miR-122

To investigate whether the significant reduction in C3U RNA abundance in the presence of miR-122 is a result of diminished RNA stability, Huh7.5 cells were transfected with HCV RNA for three days and subsequently treated with the nucleoside analog MK-0608 to block viral RNA synthesis. Total RNA was isolated at the indicated time points following drug treatment and the rate of HCV RNA decay was examined by Northern blot analysis. Data showed that wild-type HCV RNA was degraded at a slightly slower rate than C3U mutant HCV RNA (Fig 4A). The approximate half-life of wild-type RNA is 3.2 hours compared to 2.6 hours for the C3U variant (Fig 4A-B). Although modest, the reduction in RNA stability was statistically significant (Fig 4C) and could potentially impact the fitness of the C3U virus.

**Fig 4.**
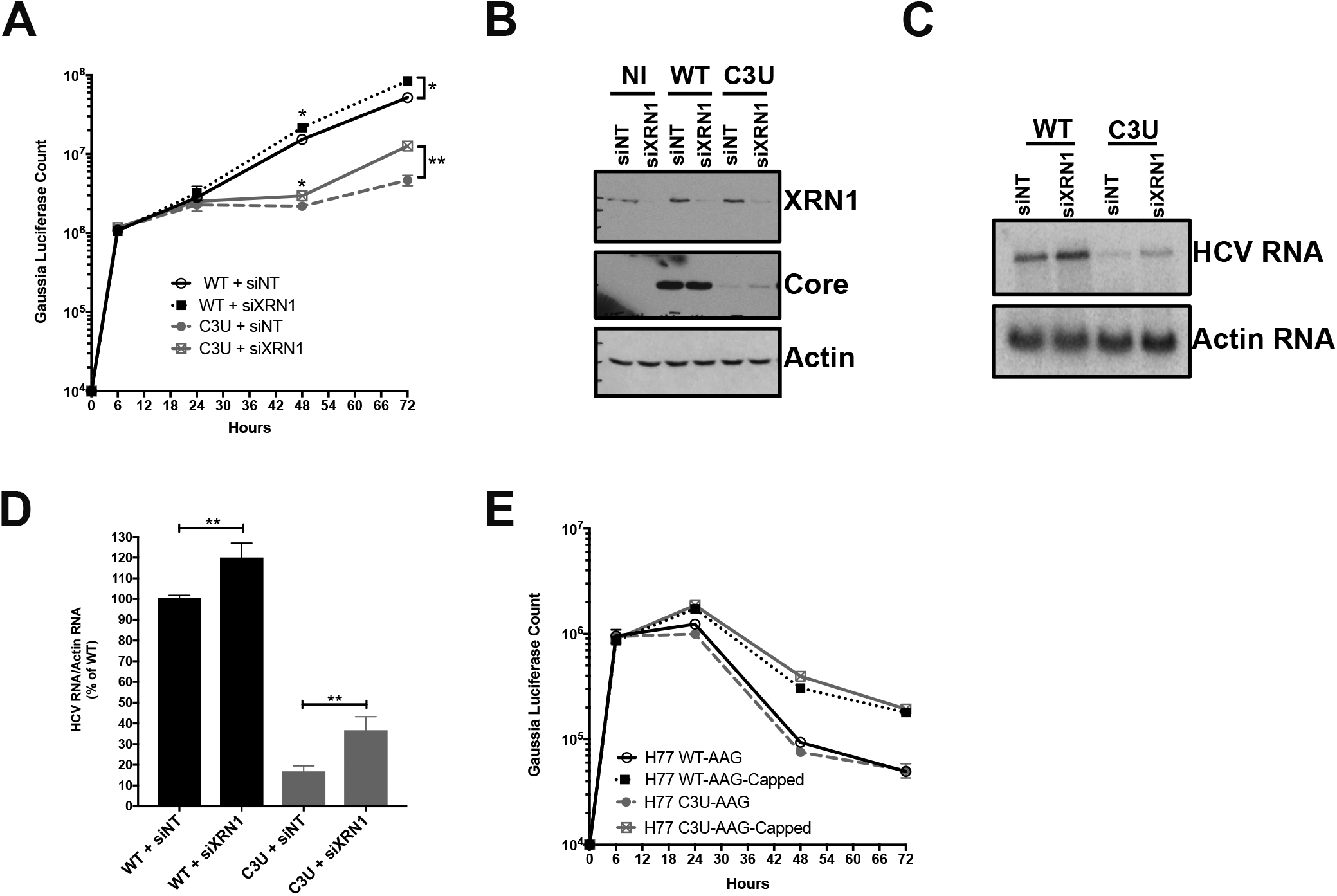
Viral RNA stability and rates of replication in wild-type and C3U variants in the presence of miR-122. (A) Effect of the C3U mutation on HCV RNA decay. Huh7.5 cells were transfected with wild-type and C3U H77.S3/GLuc RNA, treated with 25 μM of MK-0608 three days later, and viral RNA abundances were monitored at specific times after treatment, using Northern blot analyses. (B) One phase decay graph of HCV RNA (R^2^ = 0.939-0.947) was determined by normalizing HCV RNA levels to loading control actin from three independent experiments. Estimated half-lives (t_1/2_) of wild-type and C3U RNA are 3.12 hours and 2.6 hours, respectively. (C) HCV RNA half-lives (t_1/2_) of three independent experiments (**p<0.0018). (C) Rates of RNA replication in wild-type and mutant HCV variant. Three days after HCV RNA transfection, RNA was labeled with 200 μM EU for 4 and 7 hours, conjugated to biotin and subsequently isolated with streptavidin beads. RNA replication rates were determined by RT-qPCR and data is normalized to wild-type HCV four hour labeling time point (**p<0.0047, ns = non-significant).

Next, the effect of the C3U mutation on the rates of HCV RNA replication were evaluated by labeling cells with 5-EU three days post viral RNA transfection. Total RNA was isolated from cells pulsed for four and seven hours, captured and quantified as previously described. 5-EU-labeling for up to seven hours revealed a significant ~four-fold increase in the accumulation of newly-synthesized wild-type RNA compared to C3U variant RNA (Fig 4D). This impaired rate of RNA synthesis coupled with a modest, but significant decrease in RNA stability are likely sufficient to explain the reduced fitness of the C3U variant in the liver.

### Effects of XRN1 depletion on HCV C3U RNA abundance

One explanation for the reduced abundance of C3U viral RNA during low abundance of miR-122 is that reduced binding of miR-122 at site 1 could render the RNA susceptible to attack by 5’ RNA triphosphatase DUSP11 [10] and subsequently to 5’ −3’ exonucleases. Therefore, we investigated whether the low abundance of HCV C3U RNA variant of type 1a could be rescued by reducing the abundance of XRN1 by siRNA-mediated gene silencing prior to HCV RNA transfection. The effect of XRN1 depletion on HCV replication was assessed by luciferase activity and Northern blot analyses of chimeric RNAs. Robust XRN1 depletion was observed in mock- and HCV-infected samples at 72 hours after RNA transfection (Fig 5B, top panel). Depletion of XRN1 significantly stimulated the abundance of both wild-type and C3U HCV RNA at 48 and 72 hours after viral RNA transfection (Fig 5A). Similarly, increased accumulation of wild-type and C3U mutant HCV RNA was detected by Northern blot analysis in siXRN1-treated samples (Fig 5C). The effects of XRN1 depletion on wild-type HCV replication are consistent with previous observations using a similar HCV cell culture system [12]. Quantification indicated that wild-type and mutant viral RNA abundances significantly increased following XRN1 depletion by the same extent (Fig. 5D). These data suggest that both wild-type and mutant HCV RNA are similarly susceptible to XRN1 attack and that the defect in C3U RNA replication is not increased degradation due to lack of miR-122 binding at site 1.

**Fig 5.**
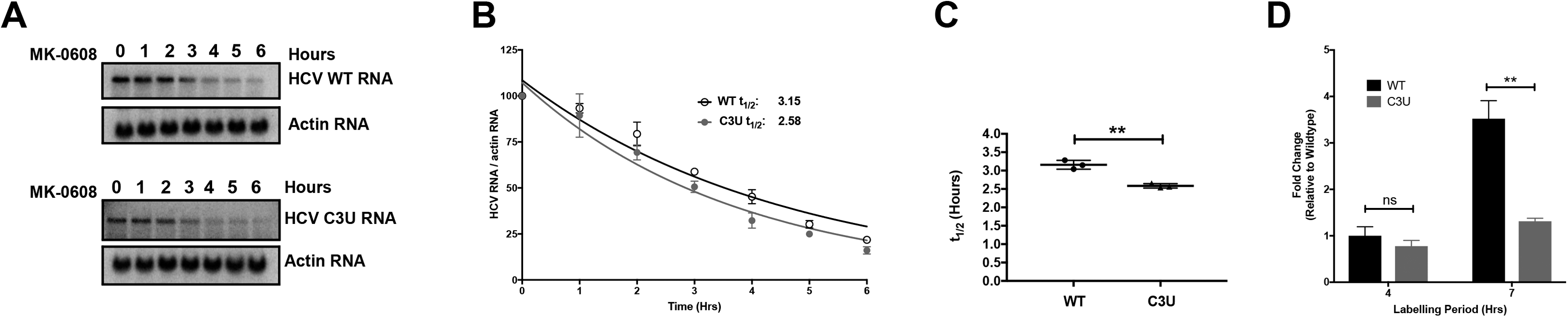
Effect of XRN1 depletion on wildt-ype and C3U HCV RNA abundances, replication, and viral protein expression. (A) Huh7.5 cells were transfected with non-targeting (NT) and XRN1 siRNAs one day prior and twenty-four hours after transfection with wild-type and C3U H77.S3/GLuc RNAs. Viral RNA replication, measured by luciferase activity in the supernatants of transfected cells, was monitored for up to 72 hours (*p<0.05 and **p<0.005). (B) Efficiency of XRN1 knockdown and its effects on HCV core viral protein levels were monitored by Western blot analysis three days post RNA transfection. (C) Cells were treated as in (A) and HCV RNA abundances were monitored by Northern blot analysis. (D) Quantification of viral RNA levels following XRN1 depletion normalized to actin. The data are representative of three independent replicates. (E) HCV RNA stability was monitored following the addition of a 5’ non-methylated guanosine cap analog to wild-type and C3U replication defective viral RNAs at the indicated time points.

To explore the possibility that differences in miR-122 occupancy in the C3U variant could alter its susceptibility to XRN1 attack further, we tested the stability of the replication defective (GDD-to-AAG) wild-type and mutant HCV RNA synthesized with and without a 5’ non-methylated guanosine cap analog after transfection into Huh7.5 cells. Viral RNAs containing a cap structure or a 5’ terminal ppp-N moiety in C3U HCV displayed similar stabilities to that of wild-type RNA across multiple infection time points (Fig 5E). Data from both lines of investigation suggest that XRN1 is unlikely to explain the observed low C3U RNA abundance in cultured liver cells.

### The C3U mutation residing in miR-122 site 1 seed sequence impairs formation of HCV:miR-122 heterotrimeric complex in vitro

The interaction between the 5’ noncoding region of HCV and miR-122 at sites 1 and 2 (Fig. 6A) extends beyond the canonical base pairing between seed and seed-match sequences [6, 7]. Mutational analysis has shown that base pairing between HCV and miR-122 at nucleotides 1-4 produces a 3’ overhang in miR-122 that shields the viral genome from subsequent exoribonuclease attack [7]. Also, previous observations indicate that the 5’ terminus of HCV forms a stable, trimolecular complex through interactions with the miR-122 at site 1 and 2 [22].

**Fig 6.**
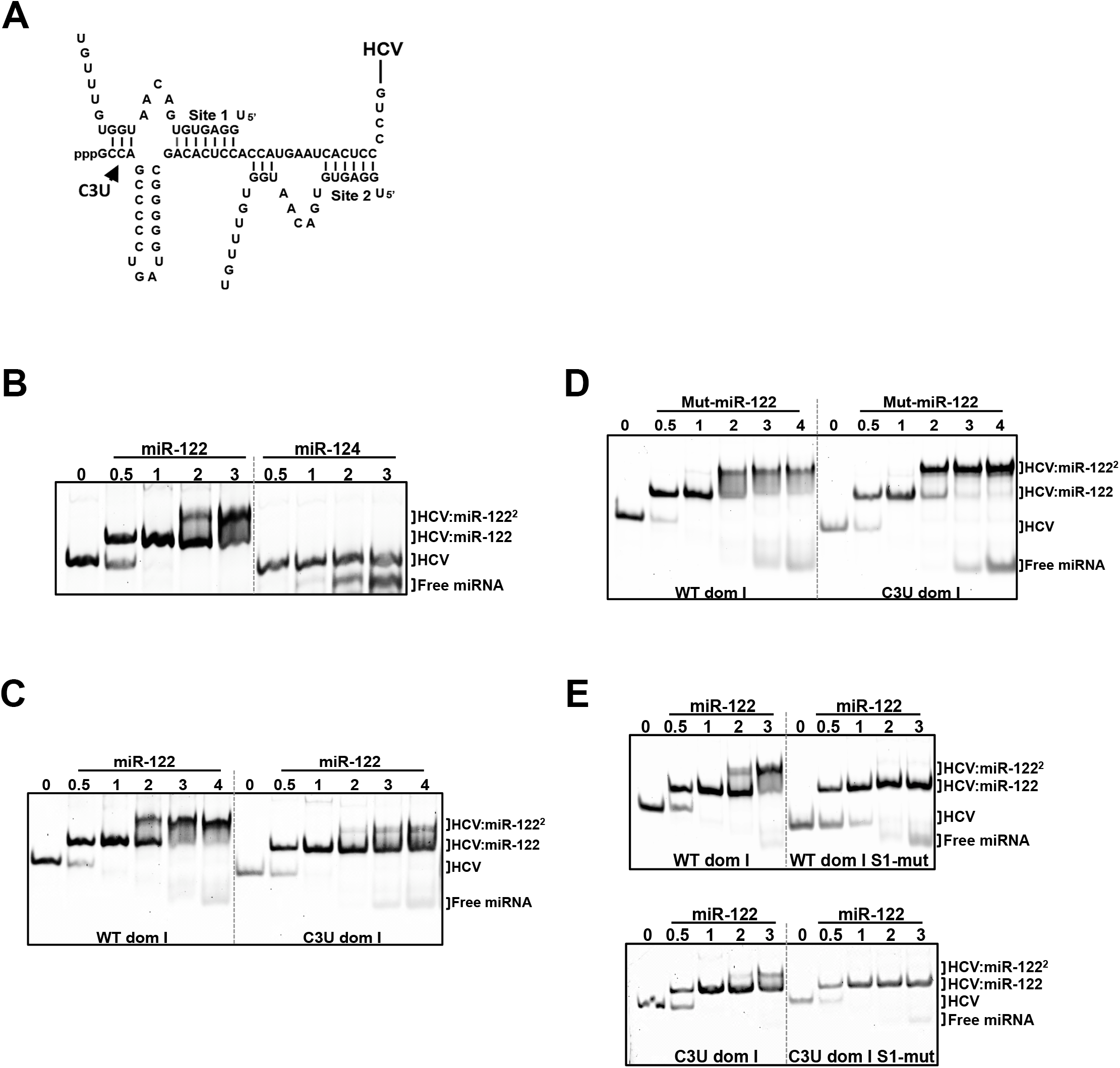
In vitro binding assays of HCV domain I nucleotides 1-47 with native and mutant miR-122 species. (A) Pairing interactions between two miR-122 molecules with 5’ RNA sequences of HCV genotype 1a (nucleotides 1-47). The position of the C3U nucleotide change is indicated. (B) HCV oligonucleotides were incubated with increasing concentrations of native miR-122 or the neuron-specific miR-124, and the resulting complexes were separated by native gel electrophoresis mobility shift assay. The migrations of free miRNA, one miR122-HCV complex (HCV:miR-122) and two miR-122-HCV complexes (HCV-miR-122^2^) are indicated. (C, D) Wild-type and C3U domain I RNAs were incubated with native miR-122 (C) or mutant miR-122 (D). (E) The seed-match sequence of miR-122 binding site 1 in wild-type and C3U RNAs was mutated and then incubated with increasing amounts of native miR-122. Resulting oligomeric complexes was resolved by non-denaturing gel electrophoresis. The data are representative of three independent replicates. Nucleic acids were visualized after staining with SYBR Gold (Thermo Fisher).

Adopting Mortimer’s and Doudna’s electrophoretic mobility shift assays (EMSA) approach [22], we investigated whether the C3U substitution disrupted the formation of the HCV:miR-122 heterotrimeric complex in solution. First, to confirm that the 5’ terminus of HCV genotype 1a (nucleotides 1-47) directly binds and forms an oligomeric complex with miR-122, wild-type HCV RNA was incubated with increasing amounts of miR-122 and the resulting complexes were resolved using EMSA. Figure 6B shows that incubation of HCV RNA with different molar equivalents of miR-122 resulted in the gradual formation of a heterotrimeric complex that migrated more slowly than free HCV RNA (Fig. 6B, lane 0). Incubation of HCV RNA with the neuron-specific miRNA, miR-124, showed no complex formation, demonstrating that the interaction between HCV and miR-122 is specific. Next, we investigated effects of the C3U mutation on the formation of the heterotrimeric complexes. Figure 6C shows that the C3U-containing RNA required a much higher concentration of miR-122 to form any heterotrimeric complex at all. To examine whether mut-miR-122 could rescue the formation of heterotrimeric complexes, wild-type or C3U RNAs were incubated with mut-miR-122 and complexes resolved by EMSA. Incubation of wild-type HCV RNA with mut-miR-122 resulted in diminished formation of the trimeric complex (Fig. 6D). In contrast, formation of a trimolecular complex was observed after incubation of C3U RNA with mut-miR-122 (Fig. 6D). These result show that mut-miR-122 can bind to both site 1 and 2 in C3U RNA, but only to high-affinity site 2 in wild-type RNA. Binding of mut-miR-122 to its target site is mediated by its interaction with the HCV RNA that extends beyond the seed-seed match interactions. To confirm that C3U RNA and miR-122 interact at binding site 2, the seed sequence of miRNA binding site 1 was mutated in both wild-type and C3U RNAs so that only site 2 was available for binding. Both RNAs did allow the formation of dimeric, but not trimeric complexes (Fig. 6E), arguing that miR-122 can bind to site 2 in C3U RNA independently of site 1. These data suggest that a single C3U mutation outside the seed-match region of site 1 results in reduced binding of miR-122 to site 1 but allows binding of miR-122 to site 2, culminating in the formation of weak, unstable trimeric complexes. It is likely that these altered miR-122/viral RNA complexes contribute to the poor viral fitness of the C3U variant in the presence of miR-122 at normal abundance.

## Discussion

Traditional antivirals have overwhelmingly focused on targeting virus-encoded proteins that are essential components of the viral life cycle using small molecule inhibitors. Due to the high mutation rates of RNA-dependent viral RNA polymerases, non-lethal mutations in the viral genome can result in drug-resistance phenotypes to these direct-acting antivirals. On the other hand, targeting the expression or function of host cellular factors that are vital for viral growth is predicted to result in a higher genetic barrier to resistance than direct-acting antivirals. The liver-expressed microRNA miR-122 was recently targeted in a miRNA-based anti-HCV therapeutic strategy. Pharmacological inhibition of miR-122 using modified anti-sense oligonucleotides (anti-miRs) significantly lowered the viral load in the serum of HCV-infected patients [17, 18].

A C3U resistance-associated mutation in the miR-122 binding site 1 of the HCV 5’ noncoding region was observed in sera from patients that experienced virologic rebound several weeks following treatment with miR-122 anti-miRs. This led us to investigate whether this single C3U substitution conferred true resistance to the inhibitory effects of anti-miRs and, if so, by what mechanism viral replication could occur at lowered abundance of miR-122. While patient-derived virus did not grow in cultured cells, using a cell culture-adapted HCV genotype 1a infectious system we found that the C3U mutation, during sequestration of miR-122, showed an increased ability to replicate the C3U RNA compared to wild-type RNA. However, this came at the expense of significantly diminished viral fitness in liver cells with normal abundance of miR-122. Specifically, RNA stability and rates of RNA replication of the C3U RNA are impaired in cells that express normal amounts of miR-122. Curiously, the C3U mutation has not been observed during selection experiments in cultured cells [17, 24, 25] and no naturally occurring C3U HCV genotypes have been deposited into Gene bank. Ono et al. observed a viral G28A mutation that arose in the serum and peripheral blood monocytes of type 2-infected patients [25], however, the C3U variant has only been observed in patient serum after several weeks in anti-miR treatment, and only transiently. Whether the extrahepatic C3U genomes reflect growth of the C3U variant in the liver of re-bounding patients is not known, because liver biopsies are not available. However, the poor fitness of the C3U variants suggests that the C3U variant may contribute little to HCV-induced liver pathogenesis. This would be similar to the situation observed with sofosbuvir, i.e. during treatment of patients with this HCV polymerase inhibitor, selection is observed for variants whose fitness in the absence of the drug is so low that they do not persist [26]

What is the mechanism by which the C3U variant can replicate in the presence of low amounts of miR-122? Using genetic and biochemical approaches, we discovered that miR-122 binds only at site 2 in the C3U RNA. Using EMSA, Mortimer and Doudna showed that miR-122 binds at miRNA binding site 2 with an affinity 50-fold greater than at site 1 [22], explaining the continued binding of miR-122 at site 2 in C3U even when miR-122 abundance is low. Why can the C3U variant accumulate in extra-hepatic cells where miR-122 abundance is very low? It is possible that novel inter- or intramolecular RNA-RNA interactions lead to stabilization of the viral genomic RNA that contains the C3U mutation in the absence of miR-122. Novel RNA-protein interactions could also take place at the 5’ end of C3U RNAs. It will be important to identify both molecular interactions that aid in its persistence, and the reservoir for the C3U variant during miR-122 anti-miR treatment.

## Materials and Methods

### Cell culture

Huh7.5 Sec14L2 and Huh7.5 ΔmiR-122 cells (generous gifts from Charles Rice, Rockefeller University, New York) were maintained in DMEM supplemented with 10% FBS, 1% non-essential amino acids, and 2 mM L-glutamine (Gibco).

### In-vitro synthesized RNA and transfection

Plasmids H77S.3 and H77S.3/GLuc genotype 1a [20] (generous gifts from Stan Lemon, University of North Carolina, North Carolina) were transcribed using the T7 MEGAscript kit (Ambion), according to the manufacturer’s instructions. Huh7.5 and Huh7.5 ΔmiR-122 cells, plated in 12-well dishes, were transfected with 1 μg of in vitro-transcribed (IVT) H77S.3/GLuc RNA using the TransIT mRNA transfection kit (Mirus Bio LLC) according to the manufacturer’s protocol. After 6 hours of incubation at 37°C, supernatants were removed for GLuc assay and replaced with fresh media. Supernatants were subsequently collected at 24 hours intervals. Supernatants were stored at -20°C before luciferase assay.

### Fluorescent focus-forming assay

Infectious titers were determined by a fluorescent focus forming units (FFU) assay. Huh7.5 cells (3×10^4^) were seeded in a 48-well plate and incubated overnight. Serial dilutions of virus stock were added to cells and incubated for six hours at 37°C. The diluted virus supernatant was removed and replaced with fresh medium. Media in each plate were exchanged daily. At day three post-infection, cells were washed once with PBS and fixed with cold methanol/acetone (1:1). HCV infection was analyzed by using a mouse monoclonal antibody directed against HCV core (Abcam) at 1:300 dilution in 1% fish gelatin/PBS at room temperature for two hours and an AlexFluor488-conjugated goat anti-mouse antibody (Invitrogen) at 1:200 dilution at room temperature for 1 hour. The fluorescent focus forming units were counted using a fluorescence microscope.

### Antisense oligonucleotides

Antisense miR-122 locked nucleic acid (LNA) and RG101 have been previously described [15, 18].

### Viral RNA quantification using a Cell-to-Ct q-RT-PCR method

3,000 Huh7.5 cells were seeded in quadruplicates in a 96-well plate one day prior to infection. The following day, cells were infected with wild-type and C3U virus at an MOI of 0.005. Six hours post infection, media was aspirated and replaced with fresh media. Media was replaced with fresh media every day. At the indicated time post-infection, cells were lysed using the Power SYBR Green Cell-to-Ct kit (Ambion), and RNAs were quantified on a Bio-rad CFX Connect quantitative-PCR (qPCR) machine and Ct values were normalized to internal control 18S ribosomal RNA expression values. The primer sequences for the HCV H77.S3 genotype were: forward 5’- CCAACTGATCAACACCAACG -3’ and reverse 5’-AGCTGGTCAACCTCTCAGGA -3’. The primer sequences for human 18S rRNA gene were: forward 5’- AGAAACGGCTACCACATCCA -3’ and reverse 5’-CACCAGACTTGCCCTCCA -3’.

### Exogenous miR-122 supplementation assay

Huh7.5 ΔmiR-122 cells were plated in 12-well dishes and transfected with annealed native miR-122 and mut-miR-122 duplexes (50nM) alone or in combination using Dharmafect I (Dharmacon) following the manufacturer’s instruction. The following day, cells were transfected with 1 μg H77 GLuc IVT RNA as stated above. Twenty-four hours post H77 RNA transfection, cells were re-supplemented with 50nM of native or mut-miR-122. Supernatants from transfected cells were collected at the indicated time points. The following oligonucleotides were used in this study: native miR-122, 5’- UGGAGUGUGACAAUGGUGUUUGU-3’; mut-miR-122, 5’-UGGAGUGUGACAAUAGUGUUUGU-3’; hsa-miR-106b, 5’-UAAAGUGCUGACAGUGCAGAU-3’.

### Nucleoside analogue MK-0608 treatment

Huh7.5 cells, plated in 60mm dishes, were transfected with 2 μg of H77 GLuc IVT RNA as stated above. Three days post-transfection, media was removed and replaced with media containing MK-0608 at a final concentration of 25 μM. RNA was collected at the indicated time points and HCV RNA was analyzed by Northern blot analysis. RNA half-lives were calculated from three independent experiments using GraphPad Prism.

### In vitro RNA capping reactions

Fifty micrograms of IVT H77 WT GLuc-AAG and H77 C3U GLuc-AAG RNAs were capped using the ScriptGap m7G Capping System (Cellscript C-SCC30610) according to the manufacturer’s protocol. RNA was extracted using the RNeasy Mini kit (QIAGEN) following the manufacturer’s instructions. Purified RNA was transfected into Huh7.5 cells and samples were collected at the indicated time points.

### Luciferase activity assays

Following RNA transfection, secreted GLuc activity was measured in 20 μl aliquots from supernatants using the Luciferase Assay System (Promega), according to the manufacturer’s instructions. The luminescent readings were taken using Glomax 20/20 luminometer using a 10 second integration time.

### Generation of H77S.3 virus

Virus production was done as previously described [27]

### H77S.3 and H77S.3/GLuc mutant generation

Nucleotide substitutions to pH77S.3 and pH77S.3/GLuc were completed using the QuickChange Site Directed Mutagenesis Kit (Agilent), according to the manufacturer’s protocol. For the C3U mutation, the following primers were utilized: 5’-ACGACTCACTATAGCTAGCCCCCTGATGGG-3’ and 5’-CCCATCAGGGGGCTAGCTATAGTGAGTCGT-3’. To introduce the lethal mutation to the NS5B polymerase (GDD>AAG), the following primer were used: 5’-CCATGCTCGTGTGTGCCGCCGGCTTAGTCGTTATCTG-3’ and 5’-CAGATAACGACTAAGCCGGCGGCACACACGAGCATGG-3’.

### Small interfering RNA transfections

For XRN1-mediated depletion, the following RNAs were utilized: sense 5’-GAGGUGUUGUUUCGCAUUAUUdTdT-3’ and antisense 5’-AATAATGCGAAACAACACCTCdTdT-3’. Sense and antisense strands were combined in 1X siRNA Buffer (Dharmacon) at a final concentration of 20 μM, denatured for 2 minutes at 98°C, and annealed for 1 hour at 37°C. As a negative control siRNA, the following oligonucleotides were used: sense 5′- GAUCAUACGUGCGAUCAGAdTdT-3’ and antisense 5’-UCUGAUCGCACGUAUGAUCdTdT-3’.

Huh7.5 cells were seeded overnight in 12 well plates. The following day, 50 nM of siRNA duplexes were transfected using Dharmafect I (Dharmacon). Following overnight incubation at 37°C, cells were transfected with H77S.3/GLuc, supernatants were collected at the indicated time points and total RNA was extracted 3 days post-transfection. Depletion of XRN1 was assessed by western blot analysis.

### Western blot analysis

Cells transfected for 3 days were washed with PBS once and lysed in RIPA buffer (50 mM Tris (pH 8.0), 150 mM NaCl, 0.5% sodium deoxycholate, 0.1% SDS, and 1% Triton X-100) in the presence of cOmplete™, EDTA-free protease inhibitor cocktail (Roche) for 30 min on ice. Lysates were clarified to remove the non-soluble fraction by centrifugation at 14,000 rpm for 10 min at 4°C. Protein concentrations were measured by Bradford Assay and 50 μg of total protein lysate was mixed with 5X protein sample buffer containing reducing agent. Samples were separated on a 10% SDS-polyacrylamide gel, transferred to a PVDF membrane (Millipore), and blocked with 5% non-fat milk in PBS-T. The following primary antibodies were used to probe the membranes: anti-Core (C7-50) (Abcam, ab2740), anti-Actin (Sigma), and anti-XRN1 (Bethyl Lab A300-443A.) Immunoblots were developed using Pierce ECL Western Blot Substrate (Thermo Fisher) following the manufacturer’s suggested instructions.

### RNA isolation and Northern blot analysis

Total RNA was extracted using the RNeasy Mini kit (QIAGEN). For Northern blot analysis of HCV and actin RNA, 10 μg of total RNA in RNA loading buffer (32% formamide, 1X MOPS-EDTA-Sodium acetate (MESA, Sigma), and 4.4% formaldehyde) was denatured for 10 minutes at 65°C, separated in a 1% agarose gel containing 1X MESA and 3.7% formaldehyde, transferred and UV-crosslinked to a Zeta-probe membrane (Bio-Rad) overnight. The membrane was blocked and hybridized using ExpressHyb hybridization buffer (Clontech) and α-^32^P dATPRadPrime DNA labeled probes.

### Nascent HCV RNA labeling

Nascent HCV RNA transcripts were quantified using the Click-iT Nascent RNA Capture Kit (Thermo Fisher) following the manufacturer’s instructions. Complementary DNA was synthesized using Superscript III reverse transcriptase (Thermo Fisher) following the manufacturer’s protocol. Newly synthesized HCV transcripts were quantified using Power SYBR Green PCR Master Mix (Thermo Fisher). HCV transcript abundances were determined by comparisons to standard curves obtained from in vitro transcribed H77.S3 RNA. The primer sequences for the HCV H77.S3 genotype were: forward 5’- CGTGTGCTGCTCAATGTCTT -3’ and reverse 5’- AATGGCTGTCCAGAACTTGC -3’. Newly C3U HCV synthesized RNA abundances were represented as fold change relative to EU labeling of wild-type HCV RNA.

### HCV-miR-122 electrophoretic mobility shift assays (EMSA)

Wild-type and mutant HCV RNA oligonucleotides corresponding to H77S.3 domain I (nucleotides 1-47) were synthesized (Stanford PAN facility). HCV domain I RNA (5 μM) was mixed with 100 mM HEPES (pH 7.5), 100 mM KCl, and 5mM MgCl_2_ in a 5 μl reaction. Reactions were heated to 98°C for 3 min, cooled to 35°C for 5 min, and incubated with miR-122 oligos in molar ratios of 0.5, 1, 2, 3, and 4 at 37°C for 30 min. For anti-miR experiments, 1 μl of miR-122 antisense oligonucleotide was added to each reaction and incubated for an additional 10 minutes. Five μl of RNA loading dye (30% glycerol, 0.5X TBE and 6mM MgCl_2_) was added and samples were separated in a non-denaturing gel (12% 29:1 acrylamide:bisacrylamide, 0.5X TBE and 6mM MgCl_2_) at 4°C for 2.5 hours at 20 Amps. Gel was stained with SYBR Gold (Invitrogen) to visualize the HCV RNA-miR-122 interactions.

### Statistical Analysis

Statistical analyses were performed with Prism 5 (GraphPad). A two-tailed paired Student’s t-test was employed to assess significant differences between two groups. Error bars represent standard error of the mean.

## Acknowledgements

We are grateful to Karla Kirkegaard for many helpful comments. The gifts of Huh7.5 Sec14L2 and Huh7.5 ΔmiR-122 cells from Joseph Luna and Charles Rice (Rockefeller University, NY) are gratefully acknowledged. We are indebted to Stan Lemon (University of North Carolina, NC) for receiving plasmids H77S.3 and H77S.3/GLuc genotype 1a.

## Author Contributions

Conceived and designed the experiments: MM, JC and PS. Performed the experiments: MM and KM. Analyzed the data: MM and PS. Wrote the paper: MM and PS. Provided background on C3U and C3U sequence information: SN.

## Supporting information captions

**Figure S1.**
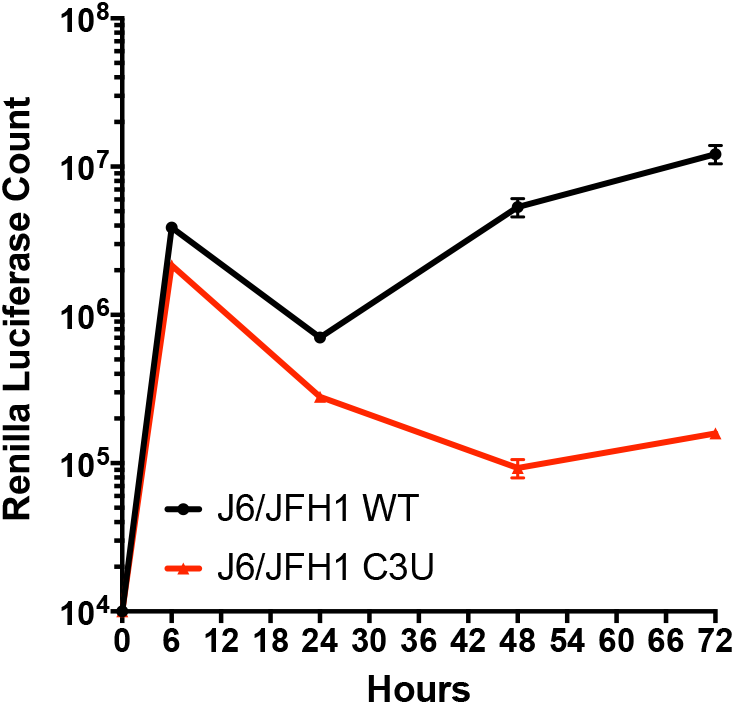
Effect of C3U mutations in J6/JHF1 viral background. Huh7.5 cells were transfected with wild-type or C3U J6/JFH1 RLuc genomes and RNA replication was determined at the indicated time points.

**Figure S2.**
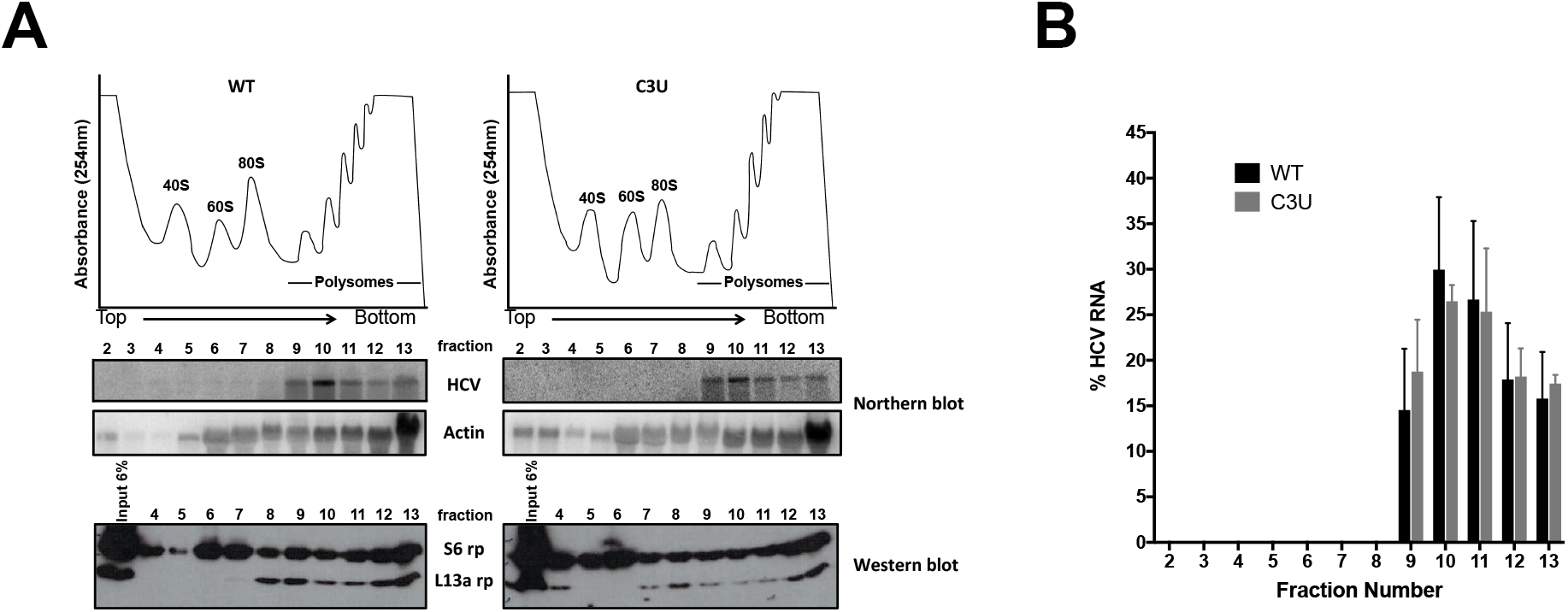
Polysomal association of C3U HCV variant. (A) HCV RNA distribution across a sucrose gradient was determined three days following transfection of Huh7.5 cells with wild-type and C3U H77.S3/GLuc RNAs. Polysomal profile trace of lysates separated in 10-60% sucrose gradients. Individual subunits, monosomal, and polysomal peaks are indicated (top). Detection of HCV and actin RNA in sucrose fractions 2 through 13 by Northern blot analysis (middle). Small (S6 rp) and large (L13a rp) ribosomal protein abundances detected by Western blot of total protein isolated from input (6%) and from fractions 4 through 13 (bottom). (B) Percent of HCV RNA distributed across the polysomal gradients of three independent experiments. Error bars display +/- SD.

## Supplemental methods

### Polysomal profiling

Huh7.5 cells were plated at a density of 5×10^6^ one day before HCV IVT RNA transfection. The following day, cells were transfected with 10 μg of HCV GLuc wild-type or C3U RNAs. Three days post transfection, 100 μg/l of cycloheximide was added to each plate. Cells were incubated for three minutes at 37°C, followed by wash with ice-cold PBS, and lysis directly on plate by adding 600 μl of polysomal lysis buffer (20mM Tris pH 7.5, 150mM NaCl, 5mM MgCl_2_, 1mM DTT, and 100 μg/mL of cycloheximide). Lysates were incubated on ice for ten minutes with periodic agitation and clarified after sedimentation at 14,000 rpm for 10 minutes at 4°C. RNA abundances in the supernatants were quantified by Nano-drop and 250 μg of cleared lysates were layered onto 10-60% sucrose gradients. Gradients were centrifuged at 35,000 rpm at 4°C in an SW41 rotor for 165 minutes and samples were fractionated with an Isco Retriever II/UA-6 detector system. RNA was extracted by adding an equal volume of acid phenol/chloroform to each fraction, centrifuged at 14,000 rpm for 15 minutes at 4°C. To remove any possible contaminants, an equal volume of chloroform was added to the aqueous fraction and centrifuged at 14,000 rpm for 10 minutes at 4°C. RNA was precipitated overnight by addition of 1 volume of isopropanol to the aqueous fraction and stored at 20°C. RNA was pelleted at 14,000 rpm for 15 minutes at 4°C, washed twice with 75% ethanol, dried for 5 minutes at room temperature, and dissolved in water. HCV and actin RNA abundances across all fractions were determined by Northern blot analysis. HCV RNA abundances per fraction were determined from three independent experiments and quantified using ImageJ.

